# Exercise intensity-dependent lactate elevation in skin interstitial fluid and its long-lasting effect

**DOI:** 10.1101/2023.10.23.563585

**Authors:** Shohei Dobashi, Daisuke Funabashi, Kazuki Sameshima, Noriko Tsuruoka, Takashi Matsui

## Abstract

Regular exercise promotes various anti-ageing adaptations in skin tissue. Although the underlying mechanisms of that might associate to the acute exercise-induced lactate signaling in the skin, it remains uncertain the profile of skin interstitial fluid (ISF) lactate dynamics during and following acute exercise. Here, we investigated whether the skin ISF lactate level increases in association with blood lactate during acute incremental exercise using a single microneedle perfusion system. The rats were acclimated to treadmill running exercise and underwent external jugular vein cannulation. Following comprehensive recovery, a 1 mm single microneedle was implanted into the back skin. Skin ISF lactate progressively increased in tandem with blood lactate during the incremental exercise but did not decrease to baseline levels until 30 minutes following the exercise unlike blood. Moreover, lactate threshold (LT), is a crucial marker of athletic aerobic performance during acute exercise, extrapolated from skin ISF showed significant alignment with blood LT. Our findings reveal that the skin ISF lactate increases associated with blood lactate and the long-lasting elevation state following acute exercise. Moreover, lactate dynamics in skin ISF can predict LT. These findings would be a milestone for elucidating regular exercise-induced physiological adaptations in the skin and evaluating athletic performance.

**Graphical Abstract:** 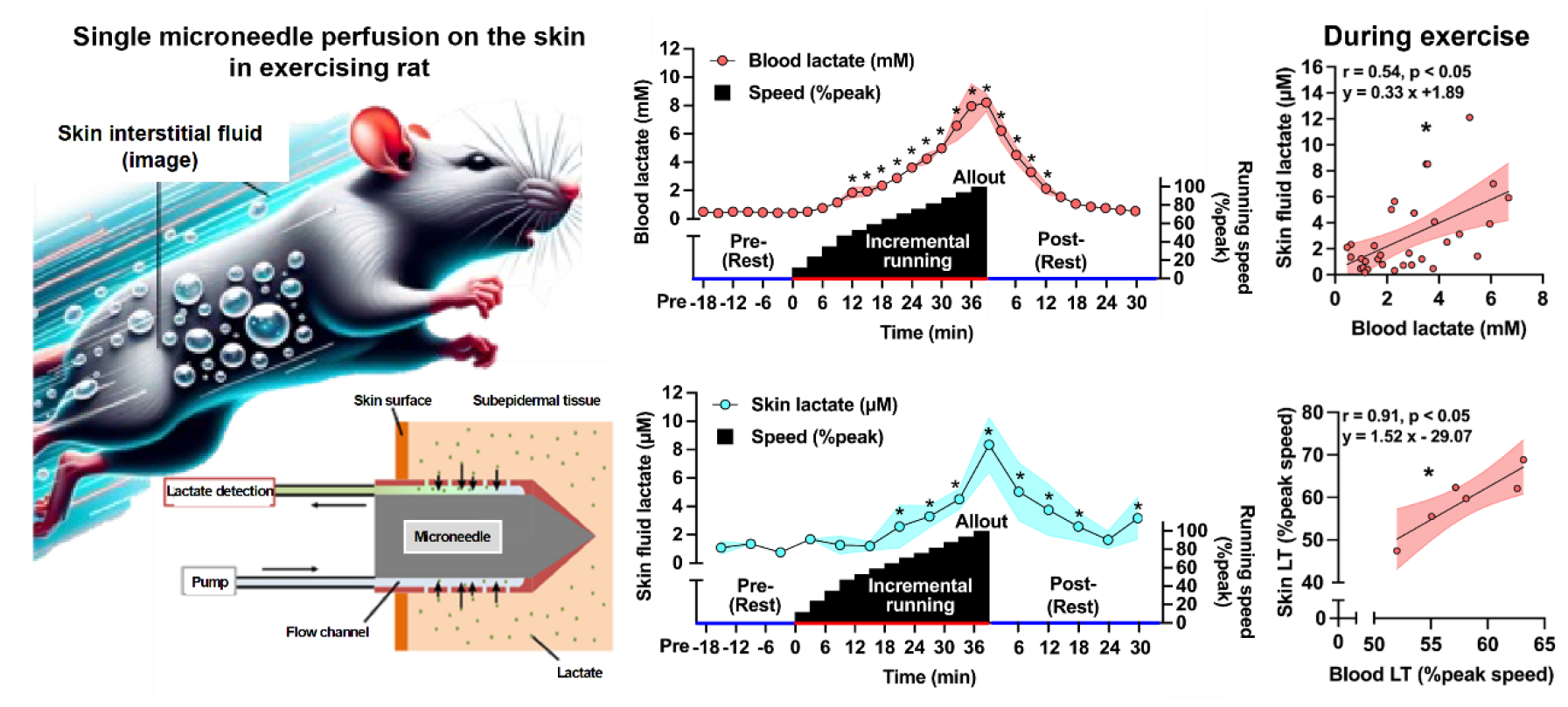

Lactate levels in rat skin interstitial fluid (ISF) increased in association with blood lactate during acute incremental exercise, but the increase in skin ISF lactate lasted 30 min following acute exercise. Lactate threshold (LT), a crucial marker of aerobic athletic performance, can be predicted from skin fluid dynamics during acute exercise. These findings would be a milestone for elucidating regular exercise-induced physiological adaptations in the skin and evaluating athletic performance.

## Introduction

The skin plays a pivotal role, such as safeguarding the body from external environmental threats, regulating body temperature, maintaining hydration, and houses sensory receptors (1); therefore, the maintenance and/or enhancement in skin tissue function are important to achieve a longer healthy lifespan. Regular exercise is believed as real polypill, which has various benefits for physical and psychological health, including skin health (2). For instance, a recent study demonstrated that regular aerobic and resistance exercises rejuvenated aging skin by reducing inflammatory factors and enhancing dermal extracellular matrices (3).

Although the underlying mechanisms of chronic exercise-induced benefits on skin health remains unknown, “lactate” is sought as one of candidates. Lactate is known not only as the end product of glycolysis, but also as an exercise-induced bioactive compound (“exerkine”) (2, 4). As the exercise intensity increases, lactate is released into the bloodstream, primarily from the exercising muscles, with a concomitant enhancement in physiological adaptations of the various tissues (5). Moreover, previous studies have reported that acute exercise-induced lactate is taken up via monocarboxylate transporters (MCTs) or binds to lactate receptors (G protein-coupled receptors, GPR81) in skeletal muscle and brain, the repetition of which results in chronic beneficial adaptations (2,6). MCTs and GPR81 are also expressed in skin (7,8) and lactate promotes mitochondrial biogenesis and attenuates inflammation there (9). Hence, the chronic exercise-induced skin benefits may also be due to the accumulation of blood-derived lactate transported to the skin, as well as to other tissues. As the constituents in blood diffuse into skin interstitial fluid (ISF) (10), we tempted to hypothesize that the skin ISF lactate level during acute exercise increases in association with blood lactate.

Recent technologies have developed to collect skin ISF and the validity of ISF lactate has been investigated (10). Among them, a novel minimally invasive microneedle perfusion system by Tsuruoka et al. appears promising, which demonstrates that the lactate dynamics in skin ISF collected using the microneedle system are consistent with those in the blood in mice after intraperitoneal lactate administration (11). Since the lactate dynamics in skin ISF were also associated with the dynamics in blood above the onset of blood lactate accumulation (OBLA) (4 mM) at this time, the microneedle perfusion system may be able to monitor dynamics of skin ISF lactate during acute exercise.

Here, we tested the hypothesis that the skin ISF lactate level increases in association with blood lactate during acute incremental exercise using the minimally invasive single microneedle perfusion system. If the results are as hypothesized, lactate dynamics in the skin ISF would allow us to predict lactate threshold (LT), which are known as universal indices that predict athletic aerobic performance and used to evaluate the exercise intensity and assessment of chronic exercise impact (12). Thus, we also aimed to examine the validity of LT determination derived from skin ISF lactate dynamics during exercise. To test this, we used rats, devoid of sweat production, because a previous study pointed out that ISF collected from the epidermal tissue may be contaminated by sweat which can interfere with adequate assessment in skin ISF lactate dynamics (13). Then, the rats were cannulated with external jugular vein and the microneedle was implanted into the back skin. This allowed for the simultaneous measurement of lactate concentrations in the blood and skin ISF during the incremental treadmill exercise.

The results of this study demonstrate that skin ISF lactate concentrations increase in association with blood lactate levels during acute incremental exercise. However, the increased skin lactate concentrations were not completely decreased to baseline levels even 30 minutes after the exercise test. We also confirmed that LT calculated from skin fluid lactate is closely related to those calculated from blood. These findings provide new insights into assessing chronic exercise-induced physiological adaptations of the skin and athletic performance using lactate dynamics in skin ISF.

## Methods

### Animals and running habituation

Adult male Wistar rats (n = 6, 250–300 g) (SLC), housed and cared for in an animal facility, were fed a standard pellet diet (MF; Oriental Yeast) and provided water ad libitum. Room temperature was maintained between 22 and 24 °C under a 12-h light–12-h dark cycle (lights on, 0700 to 1900 h). All experimental protocols were conducted in accordance with the guidelines of the University of Tsukuba Animal Experiment Committee. The rats were habituated to running on a treadmill (Natsume, Tokyo, Japan) for five sessions over 6 days for 30 min/day. The running speed was gradually increased from 5 to 25 m/min (14-16).

### Surgery to insert jugular catheter

After habituation to treadmill running, the rats were anesthetized with isoflurane and a silicone catheter was inserted into the jugular vein and fixed with a silk thread suture (32 mm). The external distal end of the catheter was fixed at the nape. Subsequent experiments and analyses were carried out 2 d after surgery (14-16).

### Inserting the minimally invasive microneedle and incremental treadmill running test

Two days after the surgery to insert catheter, the ultra-minimally invasive microneedle was inserted into the back skin of the rats under anesthetization 30 min before start of the exercise test (Figure 1*A*). This single microneedle system has an outer diameter of approximately 0.2 mm and a length of 1-2 mm can collect ISF through saline perfusion (Figure 1*B*) (11). To place the needle in the subepidermal tissue, the needle was inserted along the skin surface (Figure 1*C*). The inlet syringe was filled with saline as the perfusate. Perfusion at a flow rate of 5 μl/min was started immediately after the needle was secured to the rat’s back. This flow rate was determined considering the efficient collecting perfusate (skin ISF mixture saline) (11).

**Figure 1.**
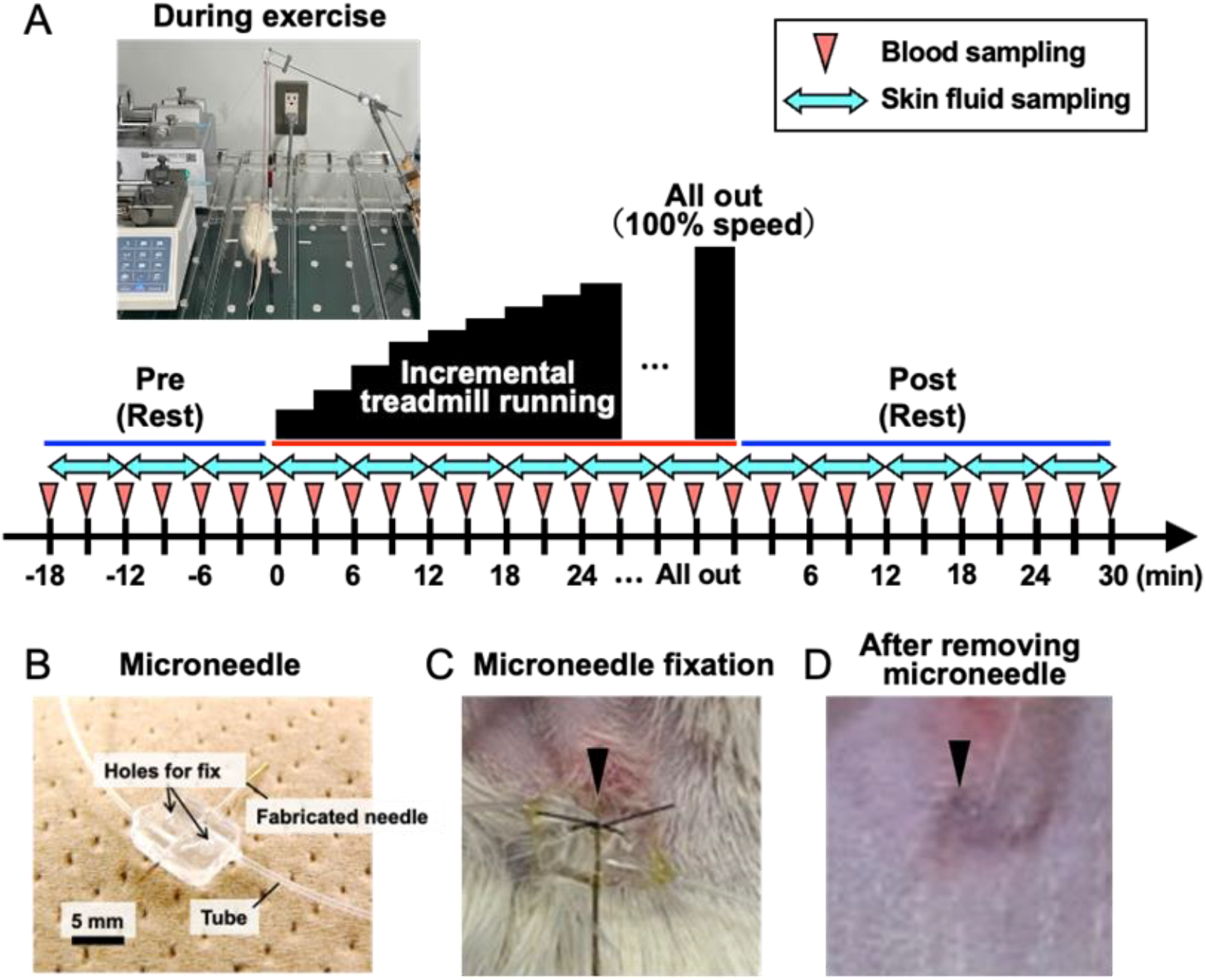
The overview of experimental procedures using a jugular catheter and an ultra-minimally microneedle perfusion system. (*A*) The experimental settings of rats with inserted catheter and microneedle perfusion system during exercise and experimental protocol of acute incremental running to determine lactate threshold for rats. (*B*) The specification of microneedle. (*C*) The part of skin with the microneedle inserted and its fixation. (*D*) The part of skin after removing microneedle. (*C* and *D*) indicate no obvious inflammation and bleeding, suggesting that the harmful effect of inserting the single microneedle perfusion system may be minimum.

After the completion of inserting microneedle into the back skin in the rats under anesthesia, the rats were moved on the treadmill and ends of the tubes were connected to syringes attached to push (ESP-36, Eicom, Kyoto, Japan) and pull (Nexus 3000, Chemyx Inc., USA) syringe pumps. 25-30 min after waking up, the animals were further kept rest for 18 min before the running test (Figure 1*A*). The initial treadmill velocity was 5 m/min and was increased 5 m/min every 3 min until the speed at 20 m/min. Thereafter, the treadmill speed was increased 2.5 m/min every 3 min until the point of exhaustion. Exhaustion was considered to have occurred when the rat was unable to keep pace with the treadmill and stayed on the grid positioned at the back of the treadmill for a period of 30 s despite being gently pushed with sticks or breathed on. This protocol is modified from an incremental treadmill exercise protocol to determine blood LT for rats (17). Following the reached exhaustion, the rats were kept rest for 35 min on the treadmill.

### Blood and skin fluid sampling

Blood samples were collected from the catheter every 3 min; however, it was collected 30 seconds before each speed increase to determine blood LT during the exercise test (Figure 1*A*). Due to the specification of the microneedle perfusion system in this study, perfusing at 5 μl/min would result in a time delay of 5 min compared to a real concentration change of skin ISF. Moreover, as the assay kit for detecting the lactate concentration needed at least 20 μl of perfusate per sample, we collected perfusate every 6 min, the collection was conducted 5 min after blood sampling from the catheter (Figure 1*A*). Theoretically, the components in the collected perfusate reflects the average values of 3 points of blood samples (5, 8, and 11 min before the skin fluid sampling). Therefore, the collected perfusate at 35 min after the exhaustion reflects until 30 min after the exercise test. In a previous study, it is suggested that the flow rate of perfusate could be changed during the recovery phase after acute exercise (18). Thus, we measured collected amount of the skin fluid in this study. We also confirmed that any fluid leaked subcutaneously or into the skin after removing the microneedle (Figure 1*D*).

### Lactate measurement and determination of lactate threshold

Immediately after each blood sampling during the exercise test, lactate levels in the whole blood were measured using an automated lactate analyzer (YSI 2500, YSI Inc., Yellow Springs, USA). Skin fluid lactate levels were determined using a commercial fluorescent stain kit (L-Lactate Assay Kit, Cayman Chemical Company, USA), according to the manufacturer’s instructions. Fluorometric determination was performed using a fluorescence microplate reader (Varioskan LUX multimode microplate reader, Thermo Fisher Scientific, USA).

We evaluated not only lactate dynamics in skin ISF with blood, but also determined lactate threshold (LT) using the modified *V-slope* method (19) in the previous study (17), calculating the running speed corresponding to the intersection of two linear regression lines derived separately from the data points of the running speed and the lactate level.

### Statistical analysis

One-way analysis of variance (ANOVA) was performed for the time course changes in blood and skin fluid lactate concentrations, and skin fluid sample amount. Pearson’s correlation analysis between the lactate concentrations in blood and skin samples was conducted separately during and following the exercise test. As mentioned above, the collected skin fluid might be reflected the 3 points of blood samples due to the methodological specification; therefore, we conducted the correlation analysis using the average concentration of 3 blood lactate per 1 skin fluid lactate (11). We also compared the absolute and relative (% peak) treadmill speed corresponding to the LT calculated from the blood and skin by paired t-test and examined their relationships between the LT calculated from blood and skin fluid by Pearson’s test. All statistical analysis was performed using the GraphPad Prism ver. 10.0.3 (MDF, Japan).

## Results

### Lactate concentration increases in the skin fluid and blood during incremental treadmill running

Following the treadmill running habituation, the rats with inserted jugular catheters and single microneedles were subjected to incremental running on the treadmill (Figure 1). The treadmill running speed was gradually increased every 3 min until exhaustion (running time: 34.4 ± 1.9 min; peak speed: 40.3 ± 1.6 m/min). Blood lactate concentrations significantly increased as running speed increased, reaching 8.2 ± 0.8 mM at exhaustion (Figure 2*A*). Skin fluid lactate concentrations gradually increased during incremental treadmill running associated with blood but did not decrease to baseline levels even 30 min after the exercise test like as blood (Figure 2*A, B*). The collected amount of the skin fluid was similar throughout the experiment (Figure 2*C*).

**Figure 2.**
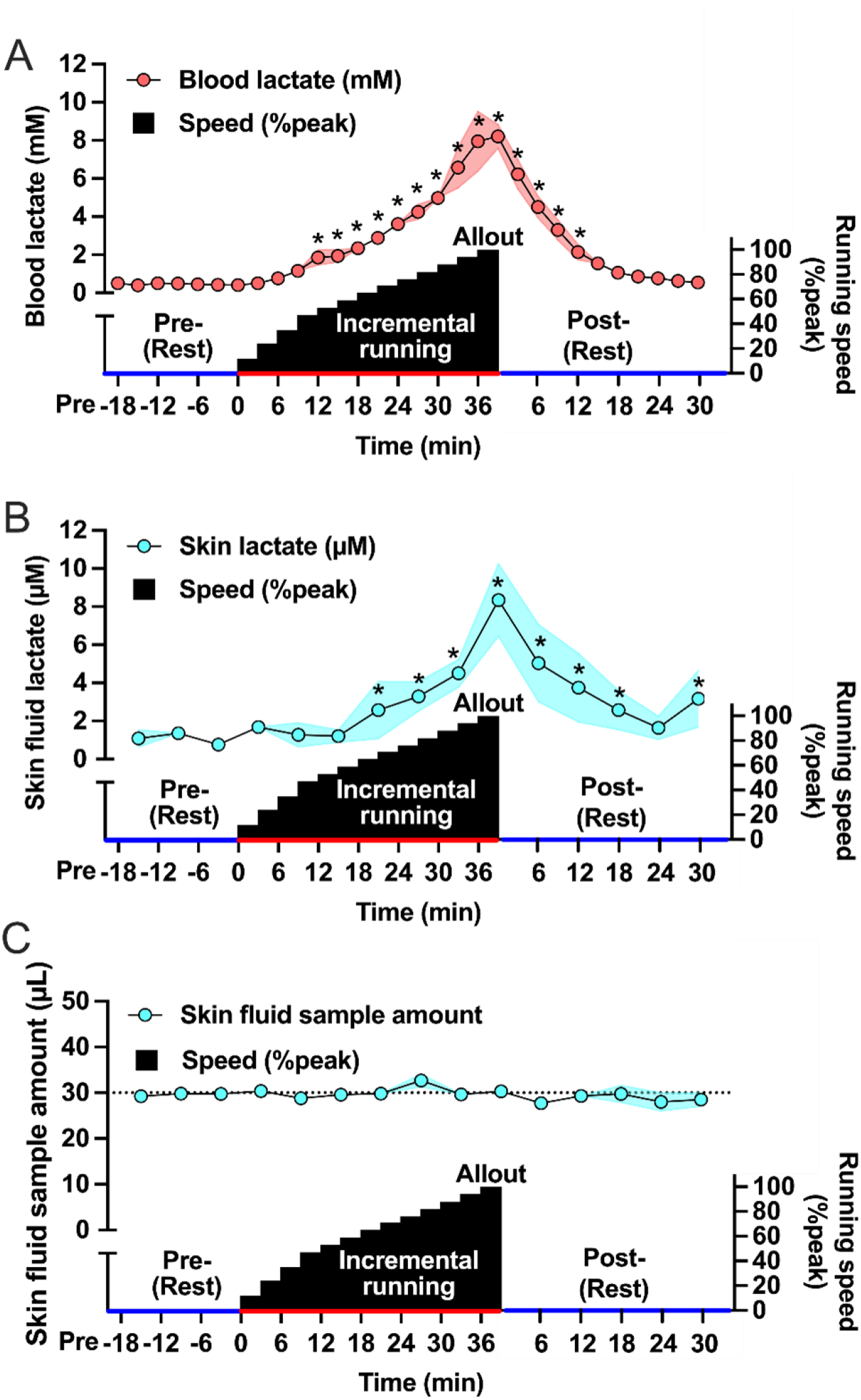
Lactate concentration increases in the skin fluid and blood during incremental treadmill running. Time course changes in blood (*A*) and skin fluid (*B*) lactate concentrations and skin fluid amount (*C*) pre, during, and post incremental running test. Each plot and shaded regions represent mean values and continuous standard error bands, respectively. *P < 0.05 vs. 0 min (immediately before the start of the exercise test).

### Skin fluid lactate dynamics are significantly associated with blood lactate dynamics during the exercise test but not afterward

We then examined the relationship between lactate dynamics in blood and skin fluid (Figure 3). A very high correlation was observed between them during exercise in individual rats (Figure 3*A, B*). The pooled analysis also revealed a moderately significant correlation coefficient (Figure 3*C*). However, these significant correlations disappeared in the recovery phase following the exercise test in both individual and pooled analyses (Figure 3*D-F*).

**Figure 3.**
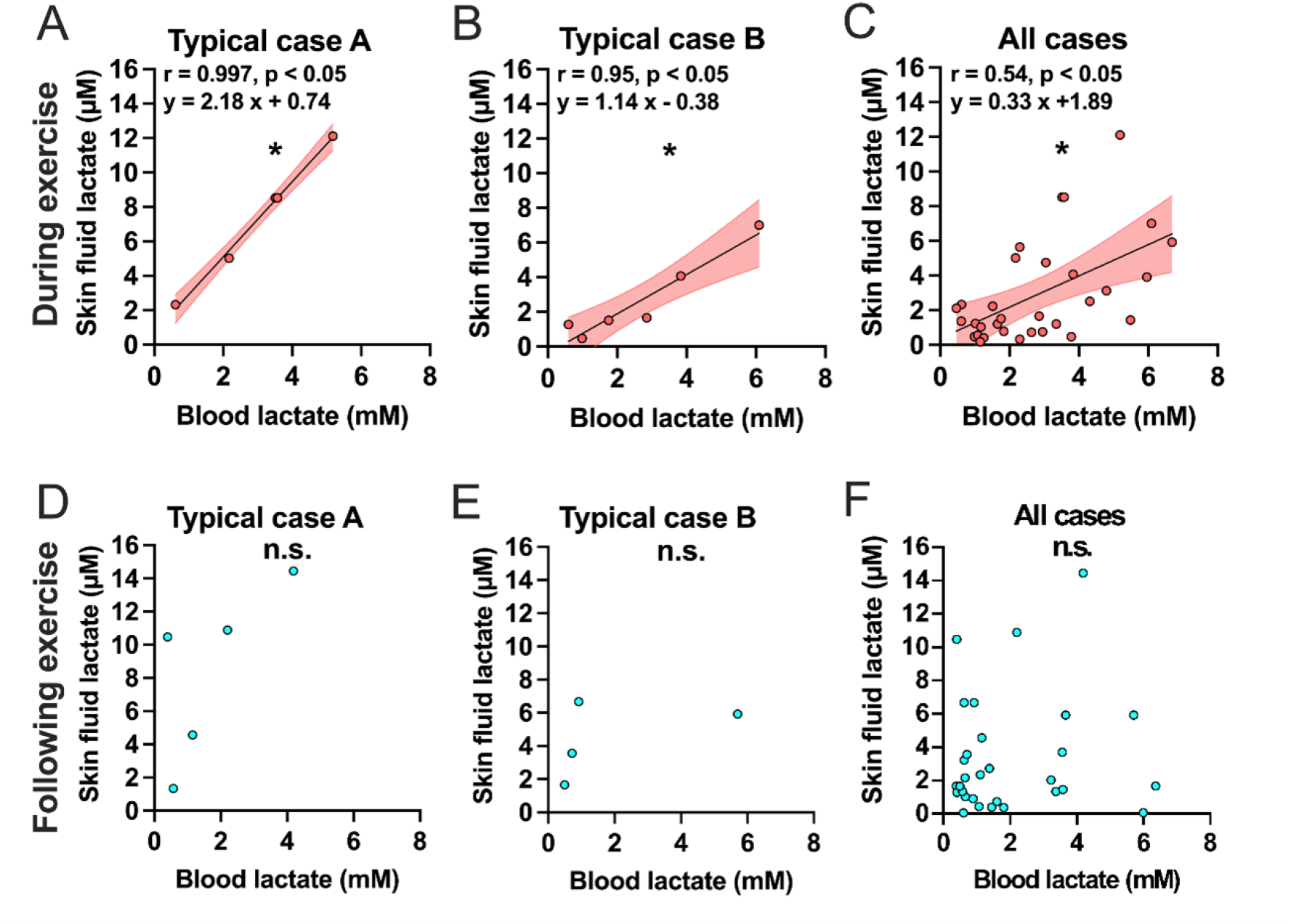
Skin fluid lactate dynamics were significantly associated with blood lactate dynamics during the exercise test but not afterward. The correlation coefficient between the blood and skin fluid lactate concentrations in typical case A and B, and pooled data of all cases during (*A-C*) and following (*D-F*) the exercise. Shaded regions represent 95% simultaneous confidence bands. *P < 0.05, n.s., not significant.

### Lactate threshold calculated from skin fluid lactate are closely related with those calculated from blood

We also attempted to calculate LT derived from skin fluid and examined its relationship with LT calculated from blood lactate dynamics by performing a correlation analysis (Figure 4). The absolute and relative (% peak) running speed and lactate levels corresponding to each LT calculated from blood and skin fluid were similar in typical participant (Figure 4*A, B*). Moreover, no significant differences were observed in the absolute and relative running speeds corresponding to the LT determined from the blood and skin fluids (Figure 4*C, E*). Furthermore, we confirmed that skin LT was significantly associated with blood LT levels (Figure 4*D, F*).

**Figure 4.**
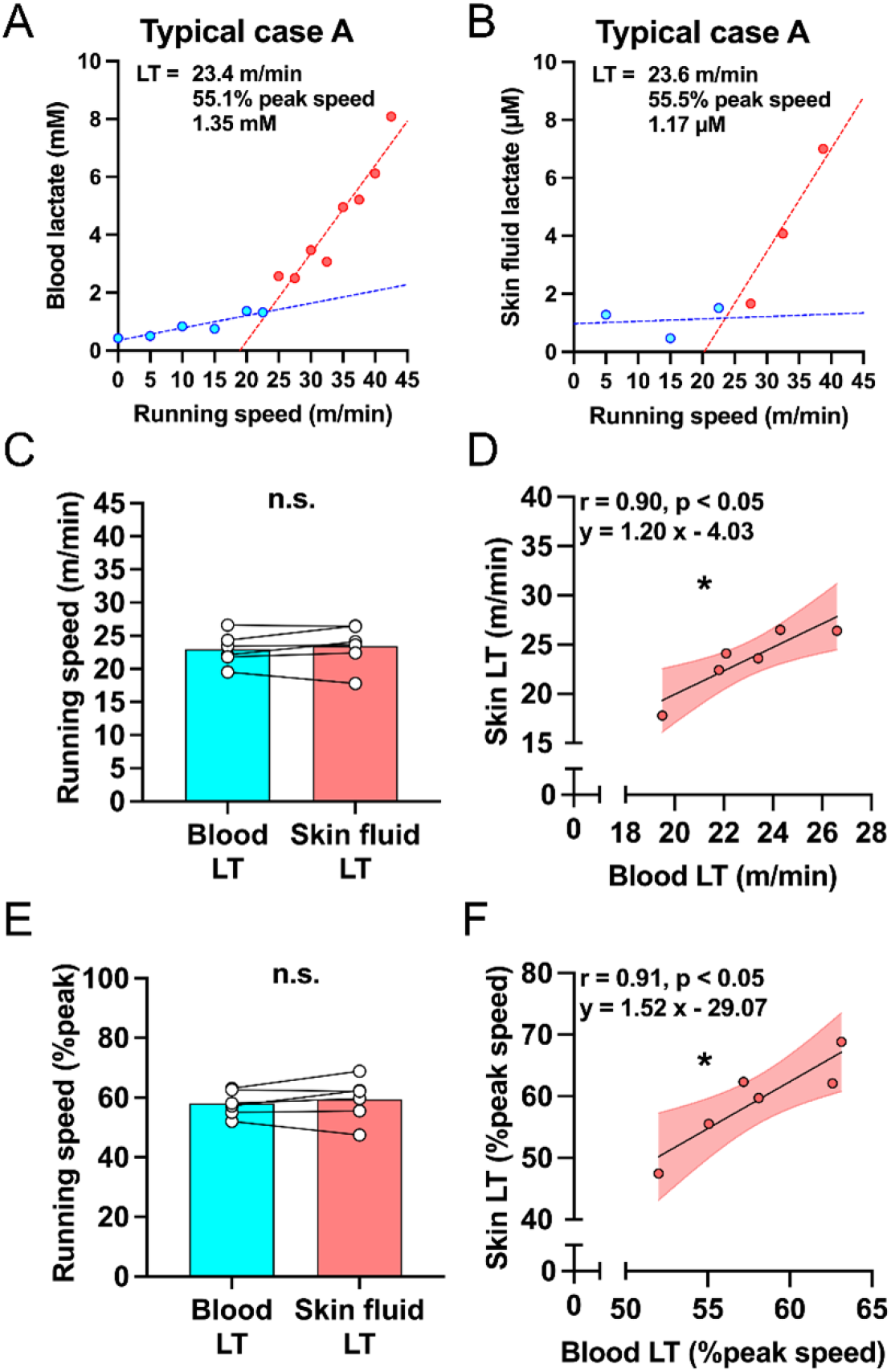
Lactate threshold calculated from skin fluid lactate are closely associated with those calculated from blood. (*A, B*) Representative LT profiles calculated by the dynamics from blood and skin fluid. The absolute running speed corresponding to the LT calculated from blood and skin fluid (*C*) and their relationship (*D*). The running speed (*E*) and its relationship (*F*) relative to peak speed during incremental exercise corresponding to LT calculated from blood and skin fluid, respectively. The values of bar graph (*C, E*) represent means and individual plots. Shaded regions represent 95% simultaneous confidence bands. n.s., not significant, *P < 0.05.

## Discussion

We tested the hypothesis that the skin ISF lactate level increases in association with blood lactate during acute exercise using the minimally invasive single microneedle perfusion system, and the results were in line with that (Figure 2). However, the increase in skin ISF lactate did not decrease to baseline levels even 30 min after the exercise like as blood lactate (Figure 2). The different linkage of lactate dynamics in blood and skin ISF during and after the exercise test was also observed in the correlation analysis (Figure 3). Additionally, we confirmed that the LT determined from the blood and skin fluids were comparable and closely related (Figure 4). As we discovered these findings in rats without sweat production in the epidermis to prevent contamination, our study provides the first evidence that unveils the skin ISF lactate dynamics response to acute exercise.

### Lactate dynamics in the perfusate from the epidermal tissue are correlated with blood during exercise

In this study, skin ISF lactate concentrations progressively increased in tandem with blood lactate during the incremental exercise (Figure 2*A, B*). As we used the rats and the lactate concentrations in perfusates collected in vitro were directly quantified by the fluorescent staining method, our present study strictly proved that skin ISF lactate concentrations itself increases as well as blood lactate. While the previous studies have pointed out that acute dynamics of lactate concentrations in ISF have delayed from the dynamics of that in blood (20), we observed that lactate dynamics in skin ISF during the acute exercise test could measure without the time delay from that in blood in this study. In this regard, a previous study demonstrated that acute exercise increased skin blood flow (21). Although further study should be required to measure skin blood flow, we speculate that the acute increase in skin blood flow due to exercise may facilitate the diffusion of water-soluble components from the blood to the skin, thereby allowing continuous lactate measurement without time delay.

Moreover, the lactate dynamics in skin fluid were significantly associated with those in blood (Figure 3*A-C*). These results are in line with those of a previous study that investigated the relationship between blood and skin fluid lactate in resting mice after intraperitoneal injection of lactate solution (11). While our study could not properly distinguish whether the skin ISF lactate was derived from the blood or the skin itself, the significant relationship between lactate dynamics in blood and skin fluids suggests that the acute exercise-induced increase in skin ISF lactate may have been derived the blood rather than in the skin. A previous study demonstrated that acute hypoxic exposure increased skin ISF lactate concentrations in association with the blood in rats (22), which supports our present findings. However, a high positive correlation of lactate dynamics during the exercise test was observed between blood and skin fluid within each individual rat (Figure 3*A-B*), the correlation coefficient of pooled analysis decreased to moderate levels (but it was still statistically significant) (Figure 3*C*). The reason for this presumably because the slope of the regression line of lactate dynamics between the blood and the skin ISF differed among the rats due to individual differences in the diffusion rate of the components from blood to skin ISF and skin blood flow rate during acute exercise. This may affect the individual variability in skin adaptation to chronic exercise. Further studies should be needed to clarify the mechanisms.

### The lactate thresholds calculated from the blood and skin fluid are closely correlated

We also observed that the LT calculated from lactate dynamics in skin ISF was corresponded to 50-65 % of peak speed as well as blood LT, and the results of the correlation analysis suggested that the individual differences were clearly captured in that range (Figure 4*F*). Although the microneedle perfusion system does not aim to estimate absolute blood lactate concentrations from skin ISF lactate concentrations, our results indicate that the lactate dynamics in skin ISF during acute exercise can predict LT. Our study used rats that conducted only running habituation without any specific training, but the relative LT was similar to that of untrained humans (approx. 50-60% peak speed) (23). Hence, the incremental exercise model in the present study seems to be a reasonable experimental model for simulating LT determination using the human incremental exercise test, and our results can be applied for human. Additionally, the exercise-intensity corresponding to the calculated skin LT is widely used in training programs for amateur-level athletes and health-enhancing exercisers (24). Steps still need to be taken toward practical application in humans, but LT assessed by skin fluid might be applicable in many sports settings in the future.

### The lactate levels in the skin fluid did not decrease to baseline levels after the exercise test

Interestingly, the increase in blood lactate concentrations rapidly decreased to baseline levels (Figure 2*A*), but those in the skin fluid decreased slowly and remained significantly higher even 30 min after the completion of the exercise test (Figure 2*C*). Additionally, the significant relationship of the lactate dynamics between the blood and skin fluid was observed during the exercise (Figure 3*A-C*) but disappeared during the recovery phase after the exercise (Figure 3*D-F*). Although it was assumed the possibility that the changes in collected amount of perfusate due to the reduction in perfusion rate of ISF following acute exercise affected lactate concentrations in the skin ISF, we observed that almost the same amount of skin perfusate was collected throughout the experiment (Figure 2*C*). This result suggests that clearance of lactate in the skin tissue may have been slower than that in tissues (mainly skeletal muscle) and the blood rather than the reduction in perfusion rate of ISF. This explanation should be supported by a rapid post-exercise decrease in skin blood flow (21), which affects the clearance of water-soluble substances from the skin into the blood (10). In any case, our results show a unique response in skin, where high concentrations of lactate last for a long-time following acute exercise.

### New hypotheses for lactate-mediated regular exercise-induced skin physiological adaptation

The present results of skin ISF lactate dynamics in this study can provide new hypotheses. Growing evidence suggests that chronic exercise improved skin conditions (2,3), and lactate acts as a signaling molecule that interacts with mitochondrial biogenesis and turnover and tunes the ability of cells to respond to stressors (9). These findings and the present results suggest that the beneficial physiological adaptation of skin tissue to chronic exercise might be due to the accumulated effects of acute exercise-induced long-lasting lactate elevation in the skin ISF. It has been demonstrated that regular higher-intensity exercise to increase blood lactate production elicited beneficial adaptations in the skeletal muscles and brain (4,5). Furthermore, the previous study that reported improved skin conditions with chronic resistance exercise also employed an intensity of exercise to increase blood lactate levels (3,25). From the fact that the lactate levels in the skin ISF increased in tandem with the blood during incremental exercise and they were closely related, regular higher-intensity exercise can be effective for inducing chronic exercise-induced skin adaptations as well as other tissue of the body. The microneedle perfusion system used in this study should be a great tool to test the new hypotheses.

### Perspectives

The microneedle perfusion system used in this study required a flow rate of 5.0 μl/min in the tubes to ensure a stable ISF amount (11). Moreover, a minimum of 20 μl of perfusate was required for the quantification of lactate concentrations by a fluorescent staining method, so the collection frequency of skin fluid had to be less than that of blood. Although our present results clearly proved that skin ISF lactate increases in tandem with blood lactate during the incremental running test, the relatively low time resolution of determination of skin ISF lactate concentrations using this system might make our interpretation difficult. In the future, a sensor that can continuously and immediately measure lactate in the perfusate should be incorporated with the microneedle to investigate the lactate dynamics in skin ISF during exercise with great precision. Moreover, the futured microneedle perfusion system should also be required to implement the technology to prevent sweat contamination for human application. The improved system will advance our understanding of lactate dynamics in skin during exercise and the resulting physiological responses. The system can also detect maximal lactate steady state (MLSS), which are used for assessing aerobic endurance capacity (26), as well as LT in real time. In this regard, we believe that our present findings to confirm the validity of LT determination derived from skin ISF dynamics will be a milestone toward realization of such innovative systems.

## Conclusion

Our findings reveal that skin ISF lactate concentrations increase in association with blood lactate levels during acute incremental exercise. However, the lactate levels in skin ISF, rather than blood, shows long-lasting elevation following acute exercise, suggesting a potential mechanism of chronic exercise-induced beneficial adaptations of the skin. Moreover, LT can be predicted from skin fluid dynamics during acute exercise without blood sampling. Collectively, we provide new insights for assessing chronic exercise-induced physiological adaptations of the skin and athletic performance using lactate dynamics in skin ISF.

## Author contributions

T.M. and N.T. conceived and designed the study. N.T. prepared a microneedle for the collection of subepidermal interstitial fluid. D.F., S.D., K.S., and T.M. collected data. D.F. and S.D. determined the blood and skin lactate concentrations. K.S. measured skin fluid amount. D.F. and T.M. conducted data analysis and interpreted the data. S.D., T.M., and D.F. drafted the manuscript. All authors edited and revised the manuscript and read and approved the final version of this manuscript.

## Acknowledgements

This research was supported in part by the Leading-edge Research Project for Sports Medicine and Science (LRP) by Japan Sports Agency to T.M., Grant-in-Aid for Scientific Research (B) by JSPS KAKENHI (19H03994 and 22H03478) to T.M., and Fusion Oriented Research for disruptive Science and Technology (FOREST) by Japan Science and Technology Agency (JST) to T.M. (JPMJFR205M) and N.T.(JPMJFR215).

## Data availability

The data that support the findings of this study are available from the corresponding author upon reasonable request.

